# Cell-type specific expression and behavioral impact of galanin and GalR1 in the locus coeruleus during opioid withdrawal

**DOI:** 10.1101/2021.01.31.428998

**Authors:** Stephanie L. Foster, Ewa Galaj, Saumya L. Karne, Sergi Ferré, David Weinshenker

**Author notes:** **Correspondence:** David Weinshenker, PhD, Department of Human Genetics, 615 Michael St, Whitehead 301, Atlanta, GA 30322.

## Abstract

The neuropeptide galanin is reported to attenuate opioid withdrawal symptoms, potentially by reducing neuronal hyperactivity in the noradrenergic locus coeruleus (LC) via galanin receptor 1 (GalR1). We evaluated this mechanism by using RNAscope *in situ* hybridization to characterize GalR1 mRNA distribution in the dorsal pons and to compare galanin and GalR1 mRNA expression in tyrosine hydroxylase-positive (TH+) LC cells at baseline and following chronic morphine or precipitated withdrawal. We then used genetically altered mouse lines and pharmacology to test whether noradrenergic galanin (NE-Gal) modulates withdrawal symptoms. RNAscope revealed that, while GalR1 signal was abundant in the dorsal pons, 80.7% of the signal was attributable to TH-neurons outside the LC. Galanin and TH mRNA were abundant in LC cells at baseline and were further increased by withdrawal, whereas low basal GalR1 mRNA expression was unaltered by chronic morphine or withdrawal. Naloxone-precipitated withdrawal symptoms in mice lacking NE-Gal (*Gal^cKO-Dbh^*) were largely similar to WT littermates, indicating that loss of NE-Gal does not exacerbate withdrawal. Complimentary experiments using NE-Gal overexpressor mice (NE-Gal OX) and systemic administration of the galanin receptor agonist galnon revealed that increasing galanin signaling also failed to alter behavioral withdrawal, while suppressing noradrenergic transmission with the alpha-2 adrenergic receptor agonist clonidine attenuated multiple symptoms. These results indicate that galanin does not acutely attenuate precipitated opioid withdrawal via an LC-specific mechanism, which has important implications for the general role of galanin in regulation of somatic and affective opioid responses and LC activity.

## 1 INTRODUCTION

Opioid withdrawal is characterized by somatic symptoms resulting from neuronal hyperactivity in multiple brain regions.^1,2^ The neuropeptide galanin is a negative regulator of neural activity,^3^ and genetic deletion of either galanin or one of its receptors, galanin receptor 1 (GalR1), exacerbates withdrawal symptoms.^4,5^ Conversely, genetic or pharmacological enhancement of galanin signaling attenuates withdrawal symptoms.^4,5^ These studies employing whole-body manipulations to galanin have prompted interest in defining the specific neuroanatomical substrates underlying the protective effects of galanin-GalR1 transmission in the brain.

Among the brain regions thought to contribute to opioid withdrawal is the noradrenergic locus coeruleus (LC),^6^ which strongly expresses galanin.^7-10^ Attenuation of opioid withdrawal by galanin is posited to involve a negative feedback loop in the LC that is maintained by galanin and GalR1.^4,11,12^ During states of LC hyperactivity such as opioid withdrawal, somatodendritic galanin release may engage Gi-coupled GalR1 autoreceptors on LC neurons, suppressing LC firing and restoring normal activity.^13,14^ Indeed, electron microscopy studies indicate that the LC is capable of dendritic galanin release,^14^ and galanin is known to induce a potent GalR1-mediated hyperpolarization of LC neurons in slice preparations.^15-19^ In addition, galanin and GalR1 are reported to be dynamically regulated in the LC by opioids, with microarray, *in situ* hybridization (ISH), and galanin reporter mouse results collectively indicating that their expression is increased by chronic morphine and precipitated withdrawal.^5,11,20^ These findings support the hypothesis that galanin and GalR1 could assuage LC hyperactive states and suppress opioid withdrawal symptoms via a local negative feedback system.

Importantly, the LC-galanin negative feedback model is predicated on the presence of both galanin and GalR1 in noradrenergic LC neurons. Galanin expression is robust in the rat, mouse, and human LC,^7-10,21^ but GalR1 expression has been difficult to characterize. Because there are no reliable antibodies for detecting GalR1 protein,^22,23^ ISH has been relied upon to examine GalR1 mRNA expression.^5,11,24,25^ Though these studies revealed the presence of GalR1 mRNA in a neuroanatomical location consistent with the LC, they lacked double labeling to confirm cellular identity and sufficiently high resolution to definitively attribute GalR1 mRNA signal to noradrenergic LC neurons. Such limitations also apply to previous studies that identified upregulation of galanin and GalR1 in the LC after precipitated withdrawal.^5,11^ Therefore, the distribution of GalR1 mRNA in the LC remains speculative, as does potential regulation of galanin and GalR1 expression by opioids.

The second tenet of the LC-galanin negative feedback model is that the LC, specifically, is the source of galanin that attenuates withdrawal severity. However, little is known about the effects of LC-derived galanin in the context of opioid withdrawal. While one study reported attenuated withdrawal symptoms in mice overexpressing galanin under the control of a noradrenergic promoter,^4^ a notable caveat is that this transgenic line exhibits ectopic galanin expression in noradrenergic neurons that do not normally contain the neuropeptide, in addition to some non-noradrenergic cells.^26^ Moreover, no study has selectively depleted noradrenergic galanin (NE-Gal) to determine whether its absence in the LC exacerbates withdrawal severity.

In this report, we sought to address remaining gaps in knowledge regarding 1) basal and opioid-induced changes in galanin and GalR1 mRNA expression in noradrenergic LC neurons, and 2) the specific role of NE-Gal in opioid withdrawal behaviors. We first employed RNAscope ISH to visualize GalR1 expression in the mouse and rat dorsal pons. We then compared GalR1 mRNA and protein expression in the mouse LC to that of adjacent non-noradrenergic neurons in dorsal pons using RNAscope and a fluorescently-tagged GalR1 transgenic mouse line, respectively. We also generated the first high-resolution, cell-type specific characterization of galanin and GalR1 mRNA expression in noradrenergic LC neurons of mice, and tested whether chronic morphine or naloxone-precipitated withdrawal altered galanin or GalR1 mRNA expression. To test whether loss of NE-Gal exacerbates withdrawal symptoms, we performed naloxone-precipitated withdrawal in genetically modified mice that lack NE-Gal (*Gal^cKO-Dbh^*). We also performed complimentary tests using NE-Gal overexpressing mice (NE-Gal OX) and WT mice treated with the galanin receptor agonist, galnon, to assess whether withdrawal symptoms could be attenuated by enhanced noradrenergic-derived or central galanin signaling, respectively.

## 2 MATERIALS AND METHODS

### 2.1 Animals

The following studies used 3-6 month old mice (both sexes) on a C57 BL/6J background unless otherwise specified. Mice were group housed on static racks with food and water available *ad libitum* in a temperature-controlled room with a 12:12 light/dark cycle unless otherwise stated. All procedures were performed during the light phase. Procedures were conducted in accordance with the National Research Council Guide for the Care and Use of Laboratory Animals and approved by the Emory University Institutional Animal Care and Use Committee.

RNAscope was performed using C57 BL/6J mice and Long-Evans rats (3 months old). Immunohistochemistry was performed using GalR1-mCherry knock-in mice, which express an mCherry tag at the C-terminus of GalR1.^27^ This strain was on a mixed 129P2/OlaHsd background and was back-crossed with C57 BL/6J mice. Withdrawal studies used previously characterized *Gal^cKO-Dbh^* and NE-Gal OX mice with respective wild-type littermates serving as controls.^10,26^ Additional details are in supplemental methods (**Methods S1**).

### 2.2 Drugs

Morphine sulfate (NIDA Drug Supply Program), naloxone hydrochloride (0.4 mg/ml stock) (Hospira, Lake Forest, IL), and clonidine hydrochloride (Sigma-Aldrich, St. Louis, MO) were dissolved or diluted in normal sterile saline. Galnon trifluoroacetate salt (Bachem, Torrance, CA) was dissolved in a vehicle of 1% DMSO in normal sterile saline. All solutions were administered using an injection volume of 10 ml/kg.

### 2.3 RNAscope

RNAscope Assay: Tissue collection, pretreatment, and RNAscope for mouse and rat LC sections was performed using the RNAscope Fluorescent Multiplex Assay v1 kit according to manufacturer’s instructions for fresh frozen tissue (Advanced Cell Diagnostics, Newark, CA). Experimental details are in supplemental methods (**Methods S1**).

Imaging: Slides were imaged using a Nikon A1R HD25 confocal microscope with NIS Elements Software. Representative images of GalR1 signal in the LC and surrounding areas were acquired with a 20x objective lens. For quantitative RNAscope studies, the LC was centered in the field of view, and a Z-stack (~14 μm thickness with 0.95 μm steps) was taken at a resolution of 1024 x 1024 pixels using a 40x objective oil immersion lens. Gain settings were chosen to maximize probe signal without oversaturation and validated with positive and negative control probe slides.

### 2.4 Immunohistochemistry

GalR1-mCherry knock-in mice^27^ were deeply anesthetized with isoflurane and transcardially perfused with 0.1M KPBS followed by 4% PFA in 0.1 M KPBS. Brains were postfixed in 4% PFA overnight and transferred to 30% sucrose in PBS for ~48 h. Brains were frozen in chilled isopentane and sectioned on a cryostat at 40 μm increments. Sections from the LC and paraventricular nucleus of the thalamus (PVT), which strongly expresses GalR1,^27^ were stained using an adapted protocol for mCherry detection in mu opioid receptor-mCherry tagged mice.^28^ See supplement for protocol (**Methods S1**). Slides were imaged using a Nikon A1R HD25 confocal microscope with NIS Elements Software. Z-stack images with 4 μm steps were acquired for LC and PVT sections after confirming lack of signal in secondary-only slides.

### 2.5 Image Analysis

All RNAscope image analysis was performed using Imaris software (Bitplane Inc., Concord, MA). Abbreviated methods are included here, with additional details in supplemental methods (**Methods S1**).

GalR1 mRNA expression in LC versus LC-adjacent neurons: To compare GalR1 mRNA expression in TH+ and TH-cells, LC sections were run with RNAscope probes for GalR1, TH, and Synaptosome Associated Protein 25 (SNAP25), a neuronal marker with an mRNA expression pattern that fills the cell soma.^29^ The image channel corresponding to SNAP25 was used to segment individual cells and associated GalR1 puncta in 3-D. The TH channel was then overlaid and used to classify SNAP25+ cells as TH+ or TH-, and the proportion of GalR1 signal was calculated by cell type. To account for differences in the number of TH+ and TH-cells observed per image, GalR1 density was also calculated. For each image, the total number of GalR1 puncta contained within a cell population (TH+ or TH-) was divided by the estimated total 2-D surface area occupied by that population in the image.

LC galanin and GalR1 mRNA regulation: LC sections from mice that received saline, chronic morphine, or underwent withdrawal were run with RNAscope probes for galanin, TH, and GalR1. Due to high fluorescent signal, galanin and TH mRNA expression were determined by acquiring respective fluorescence intensity values from each LC cell and calculating the average intensity per image. For GalR1, the same segmentation process was used as in the GalR1 SNAP25 analysis, except that the TH channel was used to segment noradrenergic LC cells and associated GalR1 puncta.

### 2.6 Naloxone-Precipitated Withdrawal

Behavior: Naloxone-precipitated withdrawal was conducted as previously described.^30^ Mice received five consecutive days of intraperitoneal (i.p.) morphine injections at 08:00 and 18:00. Morphine doses escalated by 20 mg/kg each day (i.e. 20, 40, 60, 80, 100 mg/kg). Body weights were recorded prior to each injection to detect differences in morphine-induced weight loss.^31^ On day six, mice were transported to a separate room, weighed, and given 100 mg/kg of morphine at 08:00. Two hours later, mice were removed from their home cages, injected with naloxone (1 mg/kg, s.c.), and placed into a transparent polycarbonate observation chamber. For pharmacology experiments, mice were pre-treated with galnon (2 mg/kg, i.p.) or vehicle 15 min before receiving naloxone, and clonidine (0.3 mg/kg i.p.) or saline vehicle 30 min before naloxone, consistent with previous studies.^4,32,33^ Withdrawal behaviors were video recorded for 30 min following naloxone injection, after which mice were weighed and returned to home cages. Fecal boli were counted at the end of the session; all other behaviors were scored during review of withdrawal videos.

Video Scoring: Withdrawal behaviors were scored by a blinded observer using Behavioral Observation Research Interactive Software.^34^ Behavioral scoring criteria are described in supplemental methods (**Methods S1**).

### 2.7 Galnon Feeding Test

Galnon is a non-selective galanin receptor agonist that crosses the blood-brain barrier,^35^ and i.p. administration of galnon attenuates feeding in mice and rats through actions on central galanin receptors.^36^ To validate the use of i.p. administered galnon in our withdrawal study, we measured whether galnon increased the latency of food-deprived mice to bite a food pellet or reduced their food consumption over a 30-min interval compared to vehicle-treated mice. See supplement for protocol. (**Methods S1**).

### 2.8 Statistical Analysis

Statistical analyses and graphs were generated in GraphPad Prism Version 8 (GraphPad Software, San Diego, CA). GalR1 SNAP25 data were compared by unpaired one-tailed t-test given prior observations that GalR1 was overwhelmingly higher in TH-cells. LC-specific data comparing expression at baseline, after chronic morphine, or after withdrawal, were analyzed by one-way or two-way ANOVA with Tukey’s multiple comparisons test as appropriate. For behavioral studies, weight loss during induction of morphine dependence was compared by twoway repeated measures ANOVA (time x genotype). Withdrawal data were assessed for equality of variance using a Brown-Forsythe test. Behaviors demonstrating equal variance across groups were compared using one-way ANOVA with Tukey’s multiple comparisons test; behaviors lacking equal variance were compared using a Kruskal-Wallis test with Dunn’s multiple comparisons test. Feeding test data were compared by unpaired one-tailed t-test, since effect direction (decreased feeding) was predetermined from the literature.

## 3 RESULTS

### 3.1 The LC exhibits low GalR1 mRNA expression

As no study has yet demonstrated GalR1 expression in verified noradrenergic LC neurons, we used RNAscope to visualize GalR1 mRNA expression in TH+ LC neurons in mice. Images revealed low LC GalR1 mRNA expression in a small number of cells, with comparatively higher expression in many cells adjacent (medial, lateral, and dorsal) to the LC (**Fig 1A**). The few cells that did co-express GalR1 and TH were located on the LC periphery, as opposed to the LC core (**Fig. 1B**). GalR1 signal in hypothalamus, a positive control region,^27^ showed robust signal as expected (**Fig. 1C**).

**Figure 1.**
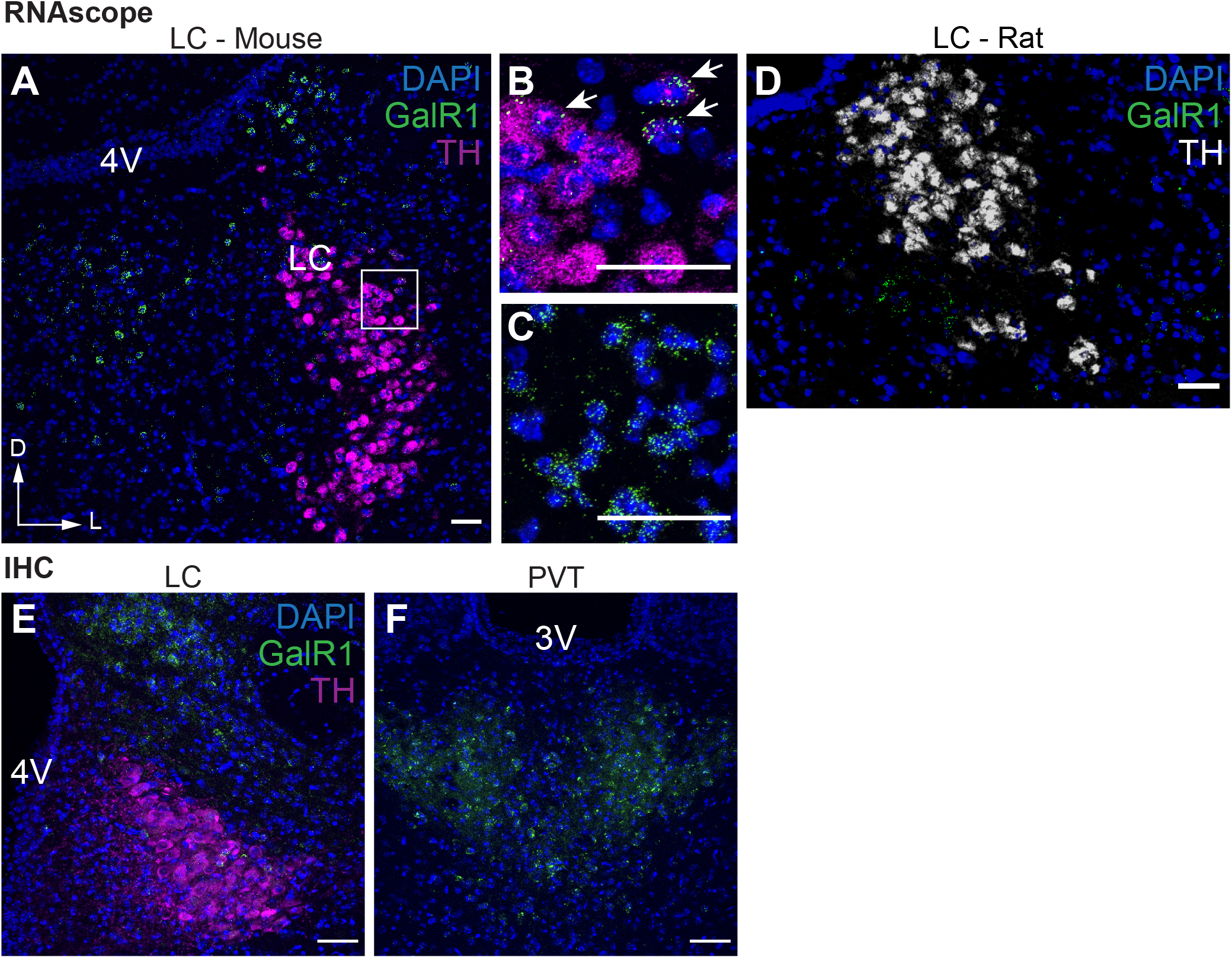
GalR1 expression is low in noradrenergic neurons of the LC. RNAscope was performed to identify GalR1 mRNA (GalR1, green) and noradrenergic neurons of the LC defined by tyrosine hydroxylase mRNA (TH, magenta) along with DAPI nuclear stain (blue). A representative image of a coronal mouse brain section shows strong GalR1 expression around, and little within, the LC (A). The few TH+ cells expressing GalR1 are observed in the LC periphery (white arrows) in comparison to the LC core (B). GalR1 mRNA was also readily observed in control sections of hypothalamus, a region previously shown to strongly express GalR1 mRNA (C). RNAscope for GalR1 mRNA (green) in the rat LC (TH, white) shows a similar pattern with GalR1 primarily outside the LC border (D). IHC for mCherry in a GalR1-mCherry mouse line reveals a pattern of GalR1 protein consistent with mRNA findings (E), and robust signal as expected in positive control sections containing the paraventricular nucleus of the thalamus (PVT, F). All scale bars are 50 μm. Abbreviations: V4 (fourth ventricle); LC (locus coeruleus); D (dorsal); L (lateral); V3 (third ventricle).

Previous work suggests that galanin attenuates withdrawal in mice but not rats.^4,37^ To assess whether this species difference is also reflected in GalR1 distribution, we performed RNAscope for GalR1 in rat LC sections. Similar to observations in the mouse, GalR1 expression was low in TH+ LC cells, yet abundant just outside this nucleus (**Fig. 1D**), demonstrating that LC GalR1 mRNA expression patterns are consistent between mouse and rat.

A potential caveat is that mRNA levels do not necessarily correlate with protein abundance. We therefore used a knock-in mouse line that expresses mCherry-tagged GalR1 protein^27^ and found that immunohistochemistry for mCherry broadly recapitulated RNAscope results. GalR1-mCherry immunoreactivity in TH+ LC neurons was negligible, but strong signal was detected just outside the LC (**Fig. 1E**), as well as in a positive control region, the paraventricular nucleus of the thalamus (PVT) ^27^ (**Fig. 1F**).

### 3.2 The majority of dorsal pontine GalR1 mRNA expression is outside of the LC

To characterize GalR1 expression in dorsal pons, we performed RNAscope using probes for GalR1, TH, and the neuronal marker SNAP25 (**Fig. 2**).^29^ The proportion of GalR1 signal was then compared between SNAP25+/TH+ LC cells and SNAP25+/TH-cells surrounding the LC. Only 19.3% of GalR1 puncta were located in TH+ cells, while 80.7% of puncta were located in TH-cells (**Fig. 2G**). An unpaired t-test of GalR1 density by cell type also indicated significantly higher GalR1 signal in the TH-cell population (*t*_10_ = 8.408, p < 0.0001) (**Fig. 2H**). These results indicate that the majority of GalR1 mRNA signal in the dorsal pons emanates from non-noradrenergic, LC-adjacent cells, rather than the LC itself.

**Figure 2.**
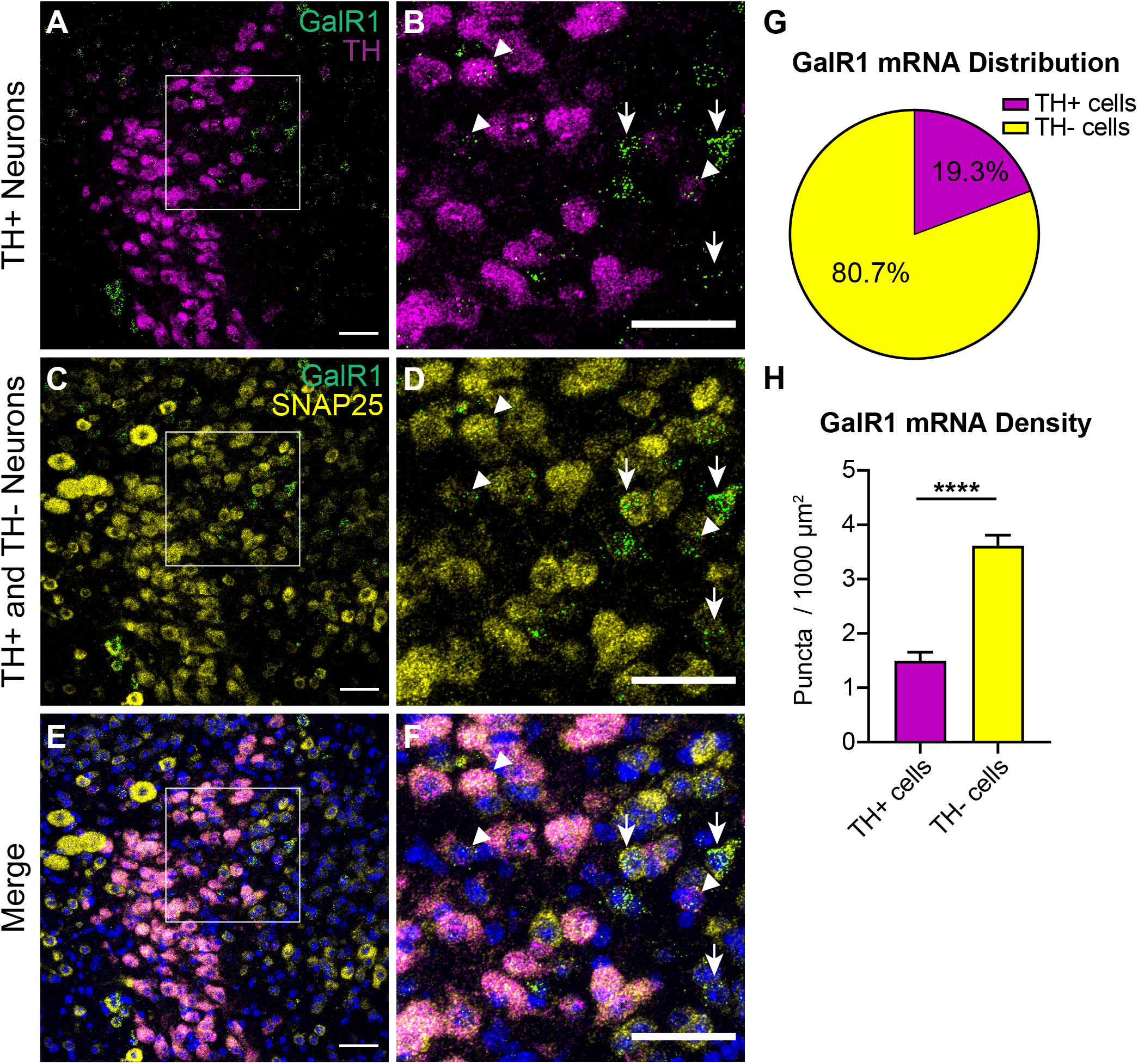
Pontine GalR1 mRNA expression is higher in LC-adjacent regions than the LC itself. RNAscope was performed for GalR1 (green) and TH (magenta) in addition to the neuronal marker SNAP25 (yellow) to evaluate GalR1 mRNA expression in TH+ and TH-populations. SNAP25 labels both TH+ LC neurons and TH-neurons in the surrounding field of view (A,C). Enlarged images show low GalR1 signal in LC neurons that are both TH+ (B, arrowheads) and SNAP25+ (D, arrowheads). Higher GalR1 signal can be seen in neurons that are TH-(B, arrows) but SNAP25+ (D, arrows). Merged images with DAPI nuclear stain (blue) highlight robust GalR1 signal in SNAP25+ neurons outside the LC (E,F). The majority of GalR1 signal in each LC image is contained within TH-, rather than TH+ cells (G). Analysis of GalR1 puncta density by cell type shows that within the LC field of view, TH-cells express significantly more GalR1 than TH+ cells (H). *n* = 6 mice, 3 LC images per mouse. All scale bars are 50 μm. Bar graphs display mean ± SEM. **** *p* < 0.0001.

### 3.3 Morphine withdrawal increases TH and galanin mRNA, but not GalR1 mRNA, in the LC

Galanin and GalR1 are reported to be dynamically regulated in the LC by opioid exposure.^5,11,20^ However, this has not been demonstrated with cell-type specificity. We therefore compared galanin and GalR1 mRNA expression in TH+ LC neurons of mice that received saline, chronic morphine, or underwent naloxone-precipitated withdrawal following chronic morphine (**Fig. 3**). TH mRNA was quantified as a positive control because its expression in LC is consistently enhanced by opioid exposure and withdrawal.^6,20,38^ For TH, a one-way ANOVA indicated a significant effect of treatment on intensity value (*F*_2,15_ = 6.739, *p* = 0.0010); intensity was significantly increased after chronic morphine (*p* = 0.0177) and withdrawal (*p* = 0.0008) compared to saline (**Fig. 3A-C; Fig. 4A**). Because TH intensity was higher in opioid-exposed groups, and TH signal was used to identify cells for LC GalR1 quantification, we wanted to ensure that these treatment groups were not biased to detect more TH+ cells than the saline group. A one-way ANOVA showed no effect of treatment group on TH+ cells detected per LC image (*F*_2, 15_ = 2.490, *p* = 0.1165), indicating that GalR1 puncta were quantified in approximately the same number of TH+ LC cells per image across treatments (saline: 102.3 ± 3.62, chronic morphine: 92.67 ± 3.64, withdrawal: 101 ± 2.55) (**Fig. 4B**). For galanin, a one-way ANOVA also showed a significant effect of treatment on intensity value (*F*_2,15_ = 6.739, *p* = 0.0082); withdrawal was significantly higher than saline (*p* = 0.0069), and there was a trend for chronic morphine (*p* = 0.0708) (saline: 200.2 ± 62.44, chronic morphine: 374.8 ± 29.74; withdrawal: 461.2 ± 55.55) (**Fig. 3 D-F; Fig. 4C**).

**Figure 3.**
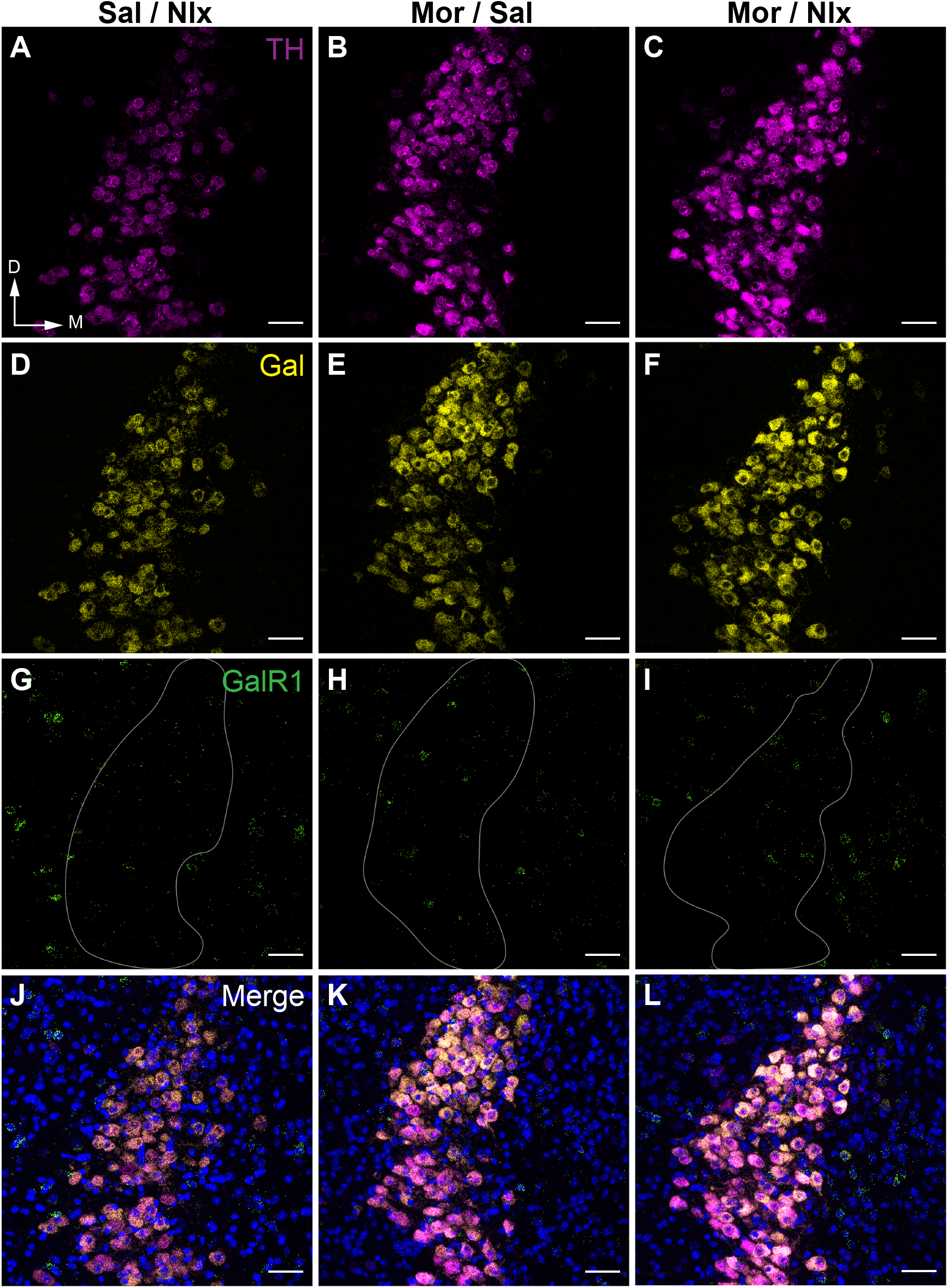
Morphine withdrawal increases LC expression of TH and galanin, but not GalR1 mRNA. Representative 40x LC images from mice that received saline injections (Sal / Nlx), chronic morphine injections (Mor / Sal), or underwent naloxone-precipitated withdrawal following induction of morphine dependence (Mor / Nlx). RNAscope was performed with probes for TH (magenta), galanin (Gal, yellow), and GalR1 (green). Compared to saline treatment, chronic morphine and withdrawal increased TH expression, indicated by elevated fluorescent signal intensity (A-C). Withdrawal also increased galanin expression compared to saline treatment (D-F). Baseline GalR1 expression was markedly lower than either TH or Gal, and expression was unaffected by chronic morphine or withdrawal (G-I, LC outlined in gray). Merged images display all probes with DAPI nuclear stain (blue) (J-L). All scale bars are 50 μm. Abbreviations: D (dorsal); M (medial).

**Figure 4.**
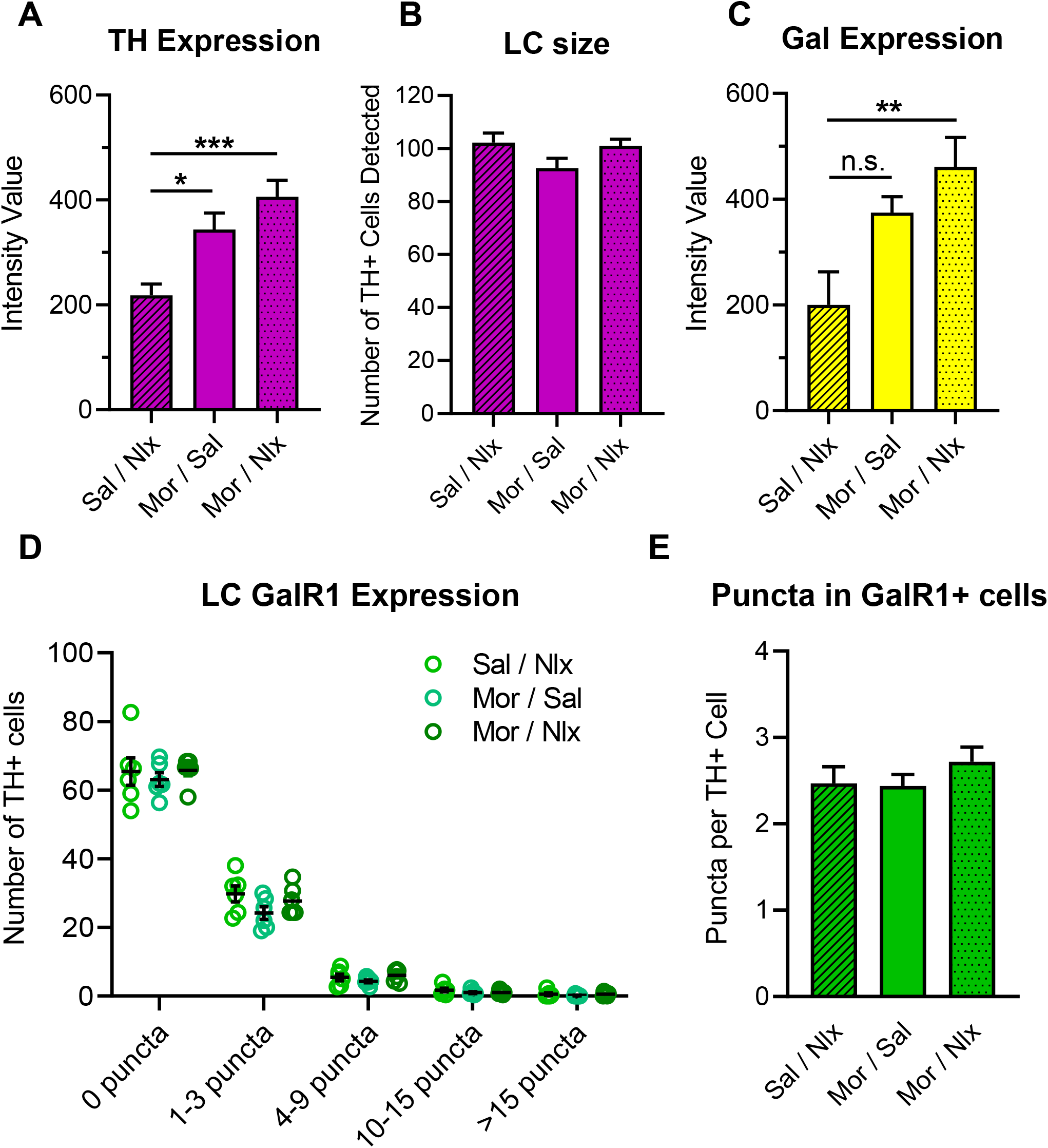
Low GalR1 mRNA expression in the LC is unaltered by chronic morphine or withdrawal. Quantification of TH, Gal, and GalR1 mRNA signal in 40x LC images from mice that received saline injections (Sal / Nlx), chronic morphine injections (Mor / Sal), or underwent withdrawal (Mor / Nlx). Withdrawal and chronic morphine increased TH mRNA expression as measured by fluorescence intensity (A). There were no differences in the number of TH+ LC cells detected per treatment group, indicating that changes in TH intensity did not affect LC quantification for GalR1 analysis (B). Withdrawal increased galanin mRNA expression (C). GalR1 is not expressed in the majority of TH+ LC neurons, and relative proportions of GalR1 expression are not altered by chronic morphine or withdrawal (D). Among cells that expressed any GalR1 puncta, treatment group did not affect the average number of GalR1 puncta per cell (E). *n* = 6 mice per group, 3 LC images per mouse. All graphs display mean ± SEM. * *p* < 0.05, \*\**p* < 0.01, \*\*\**p* < 0.001.

Due to comparatively low expression levels, GalR1 expression was analyzed by binning TH+ cells in each LC image by the number of GalR1 puncta per cell. A two-way ANOVA (bin x treatment) showed a main effect of bin (*F*_4,75_ = 929.1, *p* < 0.0001), but not treatment (*F*_2,75_ = 2.440, *p* = 0.0941), and no interaction (*F*_8,75_ = 0.5410, *p* = 0.8220), indicating that treatment did not influence the relative proportions of GalR1 expression (**Fig. 3G-I; Fig. 4D**). The majority of TH+ cells did not exhibit any GalR1 puncta (saline: 65.39 ± 4, chronic morphine: 63.06 ± 1.98, withdrawal: 65.72 ± 1.59). Approximately one third of cells contained between one and three GalR1 puncta (saline: 29.78 ± 2.31, chronic morphine: 24.17 ± 1.85, withdrawal: 27.72 ± 1.75), with the remaining small proportion containing four or more puncta. To examine GalR1 among the cells that expressed the transcript, we performed a second analysis restricted to cells containing one or more GalR1 puncta. Again, one-way ANOVA indicated no effect of treatment group on the average number of GalR1 puncta per cell (*F*_2,15_ = 0.5221, *p* = 0.6037), with similar values across treatments (saline: 2.47 ± 0.19, chronic morphine: 2.44 ± 0.14, withdrawal: 2.72 ± 0.17) (**Fig. 4E**). These results indicate that while TH and galanin mRNA increase in the LC following chronic morphine and/or withdrawal, GalR1 expression does not change from low baseline levels.

### 3.4 Noradrenergic-derived galanin does not modulate precipitated withdrawal symptoms

To determine whether selective depletion of NE-Gal exacerbates withdrawal symptoms, naloxone-precipitated withdrawal was performed in *Gal^cKO-Dbh^* mice and WT littermates alongside a cohort of NE-Gal OX mice and their WT littermates as a control for NE-Gal modulation of withdrawal symptoms.^4^ Two-way repeated measures ANOVAs showed that during the morphine dosing period, there were no differences in weight loss between *Gal^cKO-Dbh^* or NE-Gal OX mice and their respective WT littermates (**Fig. 5A, B**). For both *Gal^cKO-Dbh^* and NE-Gal OX analyses, there was a main effect of time (*Gal^cKO-Dbh^*: *F*_3.223,61.23_ = 73.72, *p* < 0.0001; NE-Gal OX: *F*_2.437,46.29_ = 97.46, *p* < 0.0001) but not genotype (*Gal^cKO-Dbh^*: *F*_1,19_ = 1.320, *p* = 0.2649; NE-Gal OX: *F*_1,19_ = 0.05025, *p* = 0.8250), and no interaction (*Gal^cKO-Dbh^*: *F*_10,190_ = 0.8423, *p* = 0.5885; NE-Gal OX: *F*_10,190_ = 0.4376, *p* = 0.9266). One-way ANOVAs for withdrawal behaviors also revealed no genotype differences for withdrawal-induced weight loss (*F*_2,39_ = 0.3357, *p* = 0.7169), jumps (*F*_2,39_ = 0.2570, *p* = 0.7746), sniffing (*F*_2,39_ = 2.119, *p* = 0.1338), paw tremor (*F*_2,39_ = 1.628, *p* = 0.2093), rearing (*F*_2,39_ = 1.028, *p* = 0.3672), wet dog shakes (*F*_2,39_ = 1.116, *p* = 0.3379), or fecal boli (*F*_2,39_ = 0.7528, *p* = 0.4778) (**Fig. 5C-G, I, J**). Only backwards steps were significantly different (*F*_2,39_ = 4. 603, *p* = 0.0160), in which *Gal^cKO-Dbh^* mice were lower than WT (*p* = 0.0124) (**Fig. 5H**). Collectively, these results indicate that neither depletion nor overexpression of NE-Gal substantially alters withdrawal behavioral profiles.

**Figure 5.**
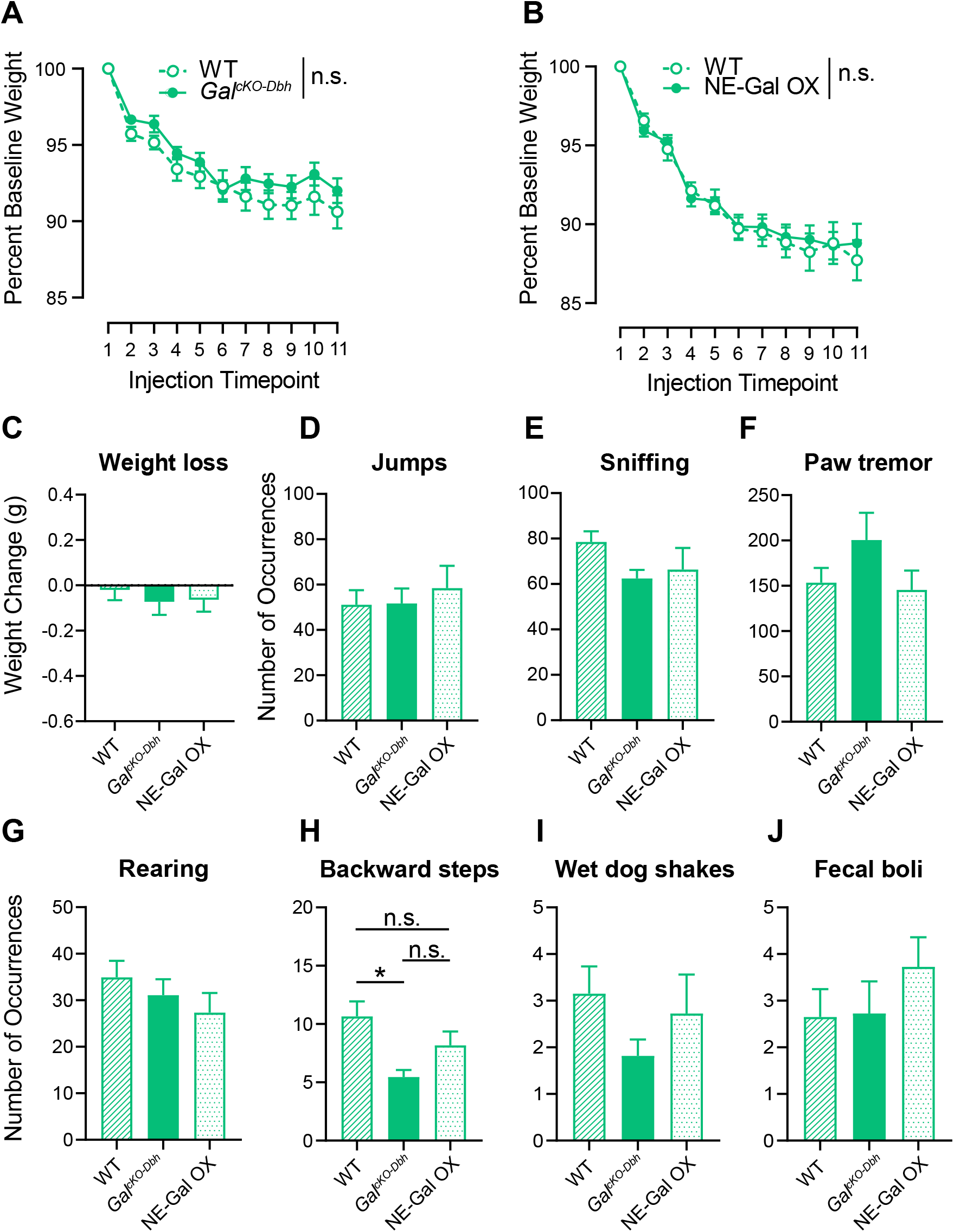
Noradrenergic galanin does not modulate precipitated withdrawal symptoms in mice. Noradrenergic-specific galanin knockout mice (*Gal^cKO-Dbh^*), noradrenergic galanin overexpressor mice (NE-Gal OX), and wild-type littermates (WT) received escalating doses of morphine twice daily for 5 days (20, 40, 60, 80, 100 mg/kg, i.p.) to induce dependence. On day 6, mice received a final dose of 100 mg/kg morphine and 2 h later underwent naloxone-precipitated withdrawal (1 mg/kg, s.c.). Neither *Gal^cKO-Dbh^* nor NE-Gal OX differed from WT littermates in weight lost during the morphine dosing period (A, B). *Gal^cKO-Dbh^*, NE-Gal OX, and WT littermates exhibited similar occurrences of most withdrawal symptoms (C-G, I, J). Only backwards steps were significantly different between groups, in which *Gal^cKO-Dbh^* exhibited fewer occurrences than WT mice (H). *n* = 10-11 for *Gal^cKO-Dbh^* and NE-Gal OX; *n* = 20 for WT. All graphs display mean ± SEM. * *p* < 0 .05, n.s. = not significant.

### 3.5 Activation of central galanin receptors does not reduce precipitated withdrawal symptoms

To confirm that peripherally administered galnon sufficiently activates central galanin receptors, we reproduced the finding that i.p. galnon reduces feeding, which requires activation of hypothalamic galanin receptors.^36^ An unpaired one-tailed t-test showed that galnon significantly reduced the amount of food consumed (*t*_12_ = 1.849, *p* = 0.0446), and increased latency to eat (vehicle 86.14 ± 38.4; galnon 179.40 ± 43.58), although this measure did not reach significance (*t*_12_ = 1.606, *p* = 0.0671) (**Fig. 6A, B**).

**Figure 6.**
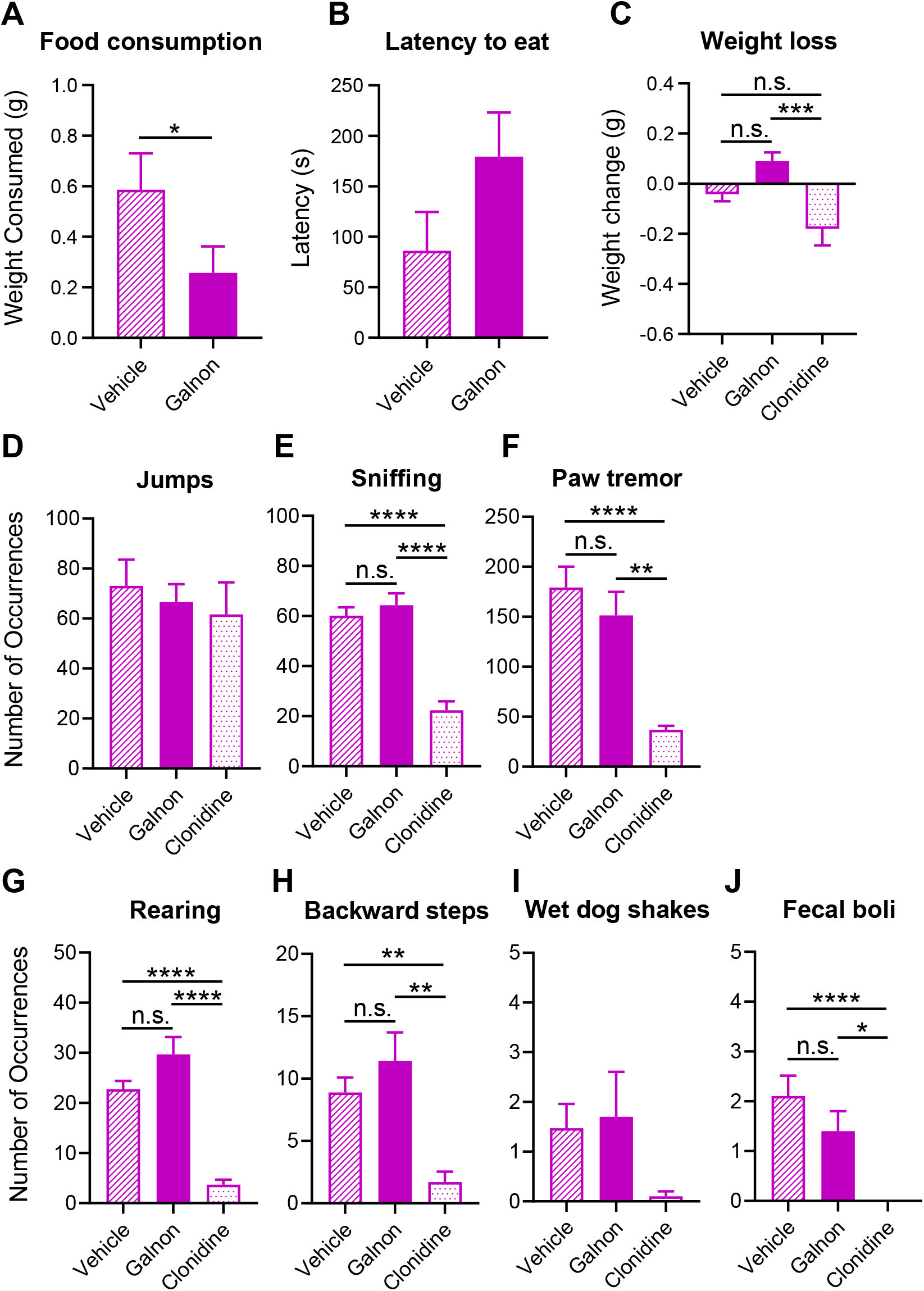
Activation of central galanin receptors does not alter withdrawal symptoms. Food-deprived mice were pre-treated with either vehicle or galnon (2 mg/kg, i.p.) prior to a feeding test. Galnon-treated mice consumed less food than vehicle treated mice (A), and latency to eat was greater than vehicle but not significantly different (B), indicating that systemic galnon was sufficient to exert central effects on feeding. *n* = 7 per group. A separate cohort was pretreated with galnon, clonidine (0.3 mg/kg, i.p.), or respective vehicle before naloxone-precipitated withdrawal. Galnon did not affect any symptoms compared to vehicle, while clonidine significantly reduced multiple symptoms compared to either vehicle or galnon treatment (E-H, J). Clonidine significantly increased weight loss compared to galnon, but not vehicle (C). Jumps and wet dog shakes did not differ from vehicle for either treatment (D, I). *n* = 10-11 for galnon and clonidine groups; *n* = 19 for vehicle. All graphs display mean ± SEM. * *p* < 0.05, ** *p* < 0.01, *** *p* < 0.001, **** *p* < 0.0001, n.s. = not significant.

We then evaluated withdrawal behaviors in mice pre-treated with vehicle, galnon, or the anti-adrenergic drug clonidine (positive control). The vehicle group included mice treated with 1% DMSO in saline (vehicle for galnon) or saline alone (vehicle for clonidine). Significant differences were detected for weight loss (*F*_2,36_ = 8.508, *p* = 0.0009), sniffing (*F*_2,35_ = 27.29, *p* < 0.0001), paw tremor (*H*_2_ = 21.55, *p* < 0.0001), rearing (*F*_2,36_ = 32.05, *p* < 0.0001), backward steps (*H*_2_ = 15.70, *p* = 0.0004), and fecal boli (*H*_2_ = 18.42, *p* < 0.0001). However, all differences were attributable to clonidine, as galnon did not reduce any symptoms (**Fig. 6C, E-H, J**). Post-hoc tests revealed that clonidine significantly reduced symptoms compared to both vehicle (sniffing *p* < 0.0001, paw tremor *p* < 0.0001, rearing *p* < 0.0001, backward steps *p* < 0.0015, fecal boli *p* < 0.0001) and galnon (sniffing *p* < 0.0001, paw tremor *p* < 0.0019, rearing *p* < 0.0001, backward steps *p* < 0.0012, fecal boli *p* < 0.0111). Unexpectedly, clonidine significantly increased weight loss compared to galnon (*p* = 0.0006), but not vehicle (*p* = 0.0538) (**Fig. 6C**). Jumps (*F*_2,36_ = 0.2856, *p* = 0.7532) and wet dog shakes (*F*_2,36_ = 1.850, *p* = 0.1719) were unaffected by either galnon or clonidine (**Fig. 6D, I**). Overall, these data imply that even broad activation of central galanin receptors fails to reduce withdrawal symptoms.

## 4 DISCUSSION

The neuropeptide galanin has been shown to modulate opioid withdrawal symptoms, which is speculated to involve an autoinhibitory feedback loop in the LC in which galanin is locally released under hyperactive conditions, and binds Gi-coupled GalR1 receptors located on LC neurons to suppress excessive activity.^4,11-13^ Surprisingly, we found that that noradrenergic LC neurons express little GalR1 mRNA, and that the majority of GalR1 mRNA signal in the dorsal pons emanates from LC-adjacent regions, rather than the LC itself. GalR1 RNAscope findings in the mouse dorsal pons were also recapitulated in rat. While precipitated withdrawal enhanced galanin mRNA expression in the LC, neither chronic morphine nor withdrawal altered GalR1 expression. Furthermore, neither decreasing nor increasing NE-Gal levels affected withdrawal symptoms. Pharmacological activation of central galanin receptors also failed to reduce withdrawal symptoms, contradicting previous reports. Our molecular and behavioral findings therefore do not support an LC-centric mechanism for galaninergic modulation of withdrawal symptoms, consistent with previous evidence that the LC is just one of several brain regions contributing to the development and expression of withdrawal.^39,40^

We found that most of the GalR1 signal in the dorsal pons, which has historically been attributed to the LC, was actually located in TH-, LC-adjacent regions including those neuroanatomically consistent with Barrington’s nucleus, the parabrachial nucleus, the pontine central gray, and the mesencephalic nucleus of the trigeminal nerve. Similar observations had been previously reported in the rat,^17,24^ but our RNAscope data now substantiate these findings with cell-type specificity, and also indicate that the pattern of GalR1 expression in the dorsal pons is conserved between rats and mice. While our study revealed basal GalR1 expression outside the LC to be higher than previously appreciated, we simultaneously found that GalR1 expression within the LC is quite low. Most unexpected was the finding that the majority of noradrenergic LC neurons do not express any GalR1, and of those cells that do, most exhibited only one to three GalR1 puncta.

These results have broader implications for the suppression of LC activity by galanin, and invite renewed discussion of previous slice electrophysiology studies. Specifically, how does galanin potently inhibit LC firing if these neurons contain so little GalR1? The first possibility is that LC GalR1 mRNA levels may not correlate with GalR1 protein; however, our data from GalR1-mCherry mice suggests that GalR1 protein is also quite low in the LC. Another explanation is that inhibitory effects of galanin are due to alternative galanin receptor subtypes, either GalR2 or GalR3. While both GalR2 and GalR3 have been identified in the rat LC,^25,41^ pharmacological and siRNA experiments do not support a GalR2-based mechanism,^15,16^ and our preliminary RNAscope results suggest that GalR3 is also sparse in TH+ LC cells of the rat (data not shown). Because the published slice electrophysiology experiments were all performed in rat, it remains unclear whether these observations pertain to mice. Alternatively, galanin may modulate LC activity through an indirect mechanism involving adjacent GalR1-rich regions that in turn affect LC firing. Previous slice electrophysiology experiments bath-applied galanin while recording from LC neurons and attributed resulting inhibition to GalR1 in the LC.^15-19^ However, our data showing that pontine GalR1 is largely outside the LC suggests that the inhibition could reflect summed actions across multiple GalR1-expressing regions, complicating interpretation of past results. The high sensitivity of RNAscope, combined with the consistent pattern of signal with our GalR1-mCherry immunohistochemistry approach, further support this possibility. Finally, it is possible that a small number of receptors may be sufficient to transduce the powerful inhibitory effect of galanin on LC firing, or perhaps many LC neurons recorded from in slice experiments happened to be those few that contained appreciable amounts of GalR1. Future studies should utilize optogenetics, cell-type specific galanin receptor knockouts, and targeted pharmacological approaches to determine whether galanin affects LC activity through a direct or indirect mechanism.

Our findings on opioid regulation of LC galanin and GalR1 expression also challenge previous work. We reproduced the finding that withdrawal enhances LC galanin mRNA expression,^5,20^ but saw no change in LC GalR1 mRNA, in contrast to a report that withdrawal increases its expression.^11^ Given that the previous ISH study lacked double-labeling, it is possible that the increased GalR1 signal included upregulation of this transcript outside the LC. However, even total GalR1 signal in our images did not differ between saline and withdrawal groups (data not shown). This disparity may be attributable to experimental differences, chiefly that we used naloxone to precipitate withdrawal, whereas the previous study used naltrexone. Additionally, we only evaluated GalR1 expression at the one significant time point used in the previous study, so there may be temporal dynamics in GalR1 expression that were not captured.

Because previous research used conventional knockout mice to argue that loss of galanin exacerbates morphine withdrawal symptoms, our data from noradrenergic-specific *Gal^cKO-Dbh^* mice do not directly refute this finding. However, the lack of altered withdrawal in *Gal^cKO-Dbh^* mice and in NE-Gal OX mice, which were reported to exhibit attenuated withdrawal symptoms,^4^ suggest that neither depletion nor enhancement of NE-Gal modulates symptom severity. Moreover, our noradrenergic-specific results are superseded by pharmacological data showing that even brain-wide activation of galanin receptors with galnon is insufficient to attenuate withdrawal, again in contrast to prior work.^4^ A caveat of our study is that galnon and clonidine were delivered systemically, so possible peripheral actions cannot be discounted. Even so, feeding test data indicate that central galanin receptors were sufficiently activated by galnon during the time frame of withdrawal evaluation, and the ability of systemic clonidine to suppress central noradrenergic transmission and opioid withdrawal is well-established.^42-45^ Our behavioral data therefore suggest that manipulations to noradrenergic and even widespread central galanin signaling do not affect withdrawal symptoms. Our finding is consistent with a report in rats in which intraventricular infusion of galanin was sufficient to modulate feeding but failed to alter naloxone-precipitated withdrawal symptoms.^37^ The lack of galanin effect reported by Holmes and colleagues was originally attributed to possible species-specific differences, or methodological limitations relating to peptide diffusion and/or proteolysis.^4^ Yet our study also failed to detect galanin effects using the same species and similar genetic and pharmacological approaches as previous mouse studies, providing an important counterpoint to the existing literature.

Although we designed our studies to align with prior work, methodological differences should be noted. Many precipitated withdrawal protocols exist, each capable of engendering different levels of symptom severity that vary widely by mouse strain.^46^ We did not use the exact protocol of Zachariou and colleagues,^4^ but both studies employed chronic, escalating morphine doses resulting in a high cumulative dose (500 versus 700 mg/kg), and the same naloxone dose (1 mg/kg). Though we evaluated withdrawal in the same NE-Gal OX mice and used C57 BL6/J mice, genetic drift can influence phenotypic differences.^47^ We also used a broader and slightly older age range for our studies (3-6 months versus 6-12 weeks). Additionally, many withdrawal studies do not include descriptions of behaviors scored, contributing to potential variation in scoring that complicates direct comparison. To that end, we provided our behavioral scoring criteria as a resource. It is possible that these collective differences impaired our ability to detect effects of galanin on opioid withdrawal; if that is the case, it would suggest that galaninergic effects are modest and require narrow experimental parameters.

One prior result we reproduced was upregulation of galanin mRNA in the LC following withdrawal,^5,20^ which implies a possible role for this source of the neuropeptide in the context of opioid use disorder, even if not detected in our withdrawal studies. Notably, the present and previous studies focused on the acute effects of galanin on somatic symptoms using precipitated withdrawal models, which are translationally similar to acute opioid detoxification in humans.^48^ But given the contribution of galanin to stress responses and depression- and anxiety-like behaviors,^3,49,50^ future work should explore whether galanin modulates affective withdrawal symptoms that arise over extended periods of time, and can be examined in spontaneous and protracted withdrawal models.^48,51^ This approach may be crucial for detecting neuropeptide effects, which develop over a longer time scale than classical fast neurotransmitters.^13^ Spontaneous withdrawal models may also more accurately reflect the human experience, in which withdrawal symptoms emerge over time due to prolonged opioid abstinence.^52^

## 5 CONCLUSION

In summary, we found that in contrast to galanin mRNA, GalR1 mRNA expression is low in the LC, and is not modulated by chronic morphine or withdrawal. Our results regarding LC GalR1 expression, in combination with behavioral data suggesting NE-Gal does not modulate withdrawal, argue against a mechanism by which acute galaninergic actions in the LC attenuate precipitated somatic withdrawal symptoms. Future work should utilize the molecular findings identified here to probe alternative mechanisms underlying GalR1 effects on LC function, as well as other behavioral aspects of opioid withdrawal.

## Acknowledgements

We thank Kathleen Smith and Laura Butkovich for technical assistance with RNAscope, L. Cameron Liles and Micah Chrenek for mouse colony management, Ellen Woon for piloting withdrawal studies, and Kevin Donaldson for assistance with image analysis. We also thank Dr. Marina Picciotto and Dr. Venetia Zachariou for their helpful input regarding this project.

## Author contribution

SLF and DW conceived, designed, and supervised the project with consultation from SF and EG. Experiments were performed by SLF and EG. SLF analyzed data with assistance from SLK. SLF and DW wrote the manuscript with input from co-authors.

