## Supplemental Methods for "Cell-type specific expression and behavioral impact of galanin and GalR1 in the locus coeruleus during opioid withdrawal"

### Animals

*$Gal^{cKO\text{-}Dbh}$* mice were generated by crossing a line expressing *cre* recombinase under the noradrenergic-specific dopamine β-hydroxylase (*Dbh*) promoter with a floxed galanin line (*$Gal^{cKO}$*) (JAX stock # 034319). *$Dbh^{cre/+}$*;*$Gal^{cKO}$* homozygotes were crossed with *$Gal^{cKO}$* homozygotes to generate *$Gal^{cKO\text{-}Dbh}$* progeny as previously described.[10] NE-Gal OX mice contain a transgene in which galanin expression is driven by the *Dbh* promoter, resulting in a five-fold increase in galanin mRNA in the LC and increased galanin immunoreactivity in LC projection regions (JAX stock # 004996).[26]

### In Situ Hybridization

Tissue Collection: Animals were deeply anesthetized with isoflurane and quickly decapitated. For RNAscope studies in mice that received saline, chronic morphine, or withdrawal, mice were sacrificed 3 h after the final injection, as previously described.[11] Brains were immediately frozen in an OCT-filled cryomold that was submerged in isopentane chilled with dry ice. OCT blocks were stored at -80°C until sectioning. Brains were sectioned at 16 μm increments onto charged slides and stored at -80°C until used for RNAscope.

RNAscope Assay: Sample pretreatment was performed as instructed using the RNAscope Sample Preparation and Pretreatment Guide for Fresh Frozen Tissue. Briefly, slides were removed from the -80°C freezer and immediately fixed in pre-chilled 10% NBF for 15 min. Slides were then dehydrated using the following ethanol wash series in 5-min increments: 50%, 70%, 100%, 100%. Slides were air dried, a hydrophobic barrier was drawn around the tissue, and slides were incubated with Pretreat IV at room temperature for 30 min. Slides were washed 2x in

PBS, experimental probe was added to each section, and slides were incubated in the HybEZ oven at 40°C for 2 h.

For qualitative images of GalR1 mRNA in mouse LC, mouse probes for GalR1 (ACD cat no. 448821), and tyrosine hydroxylase (TH, a marker of noradrenergic cells) (ACD cat no. 317621) were used. Mouse multiplex positive (ACD cat no. 320881) and multiplex negative (ACD cat no. 320871) control probes were used to validate experimental probe signal. For qualitative images of GalR1 signal in rat LC, rat probes for GalR1 (ACD cat no. 439791) and TH (ACD cat no. 314651) were used. Experiments analyzing GalR1 in TH+ and TH- neurons within the LC field of view used probes for GalR1, TH, and the neuronal marker Synaptosome Associated Protein 25 (SNAP25) (ACD cat no. 516471-C3). For the LC-specific analysis of GalR1 mRNA expression at baseline and after chronic morphine or withdrawal, mouse probes for GalR1, galanin (ACD cat no. 400961), and TH were used.

After hybridization, slides were washed 2x 2 min with wash buffer. Four subsequent rounds of amplification and 2x 2 min washes with wash buffer were performed as instructed. In amplification step 4, color module Alt A-FL was chosen to assign the following fluorophores to each channel: C1 Alexa 488, C2 Atto 550, C3 Atto 647. Slides were then coverslipped using Prolong Diamond Antifade Mountant with DAPI (Thermo Fisher Scientific, Waltham, MA) and stored in the dark at room temperature overnight. All slides were imaged between 24 to 48 hours after performing RNAscope.

**Image Analysis**

All RNAscope experiments contained 6 mice per group. For each mouse, 3 LC images were analyzed. Values were averaged across images for each mouse, and then across mice for each group.

GalR1 mRNA expression in LC versus LC-adjacent neurons: The image channel corresponding to the SNAP25 probe was isolated, and a Surface layer was generated in Imaris to identify and segment individual SNAP25+ cells in 3-D. The channel corresponding to the GalR1 probe was then used to generate a Spots layer identifying individual GalR1 puncta. The GalR1 puncta were then filtered to select only for puncta contained within the surfaces of the SNAP25+ cells identified in the Surface layer. Then, the TH channel was overlaid and used to label each SNAP25+ cell as either TH+ or TH-. GalR1 puncta per cell counts were generated for each cell, which was also classified as TH+ or TH-. Distribution of GalR1 puncta by cell type was determined for each image by dividing the total puncta within a cell population (TH+ or TH-) by the total puncta within SNAP25-defined cells. To account for differences in the number of TH+ and TH- cells observed per image, GalR1 density was also calculated by cell type. The total 2-D surface area for each cell population was calculated by summing the surface areas of the individual cells for that population within each image. Then, the total GalR1 puncta contained within a cell population was divided by the estimated total 2-D surface area occupied by that population in the image.

LC galanin and GalR1 mRNA regulation: The image channel corresponding to the TH probe was isolated, and a Surface layer was generated in Imaris to identify and segment individual TH+ cells in 3D. The channel corresponding to the GalR1 probe was then used to identify and filter GalR1 puncta as in the SNAP25 analysis.

**Immunohistochemistry**

Sections were washed in PBS 3x for 10 min each and incubated in blocking solution (PBS-Triton (0.3%), 2% normal goat serum, 1% BSA) for 1 h at room temperature, then incubated with primary antibodies overnight at 4°C. A rabbit anti-DsRed polyclonal antibody

(Takara Bio, Mountain View, CA, cat no. 632406, dilution 1:1000) was used for mCherry detection, and a chicken anti-TH polyclonal antibody (Abcam, Cambridge, MA, cat no. ab76442, dilution 1:1000) was used to define noradrenergic LC neurons. The next day, sections were washed in PBS 3x for 10 min each. Then sections were incubated in goat anti-rabbit 488 conjugated (Invitrogen Cat no. A11008, dilution 1:2000) and goat anti-chicken 633 conjugated (Invitrogen A21103, dilution 1:600) secondary antibodies for 2 h at room temperature, washed in PBS 3x for 10 min each, mounted onto charged slides, and dried overnight. Slides were coverslipped using Fluoromount-G with DAPI (Thermo Fisher Scientific, Waltham, MA, cat no. 00-4959-52) the following day.

**Withdrawal Video Scoring Criteria**

For every mouse, each occurrence of the following behaviors during the 30-min observation period was marked as a point event: rearing, jumping, wet dog shakes, paw tremor, backwards steps, and sniffing. The criteria below were used to score each behavior:

Rearing: Mouse supporting itself on extended hind legs.

Jumping: All four paws leaving the ground at the same time.

Wet dog shake: Brief, rapid shake involving both the head and body of the mouse.

Backwards steps: Mouse jumping or shuffling backwards, with visible movement of hind paws.

Paw tremor: Brief, rapid shaking of one or both front paws or hindpaws. During instances where this occurred rapidly, each "bout" of tremor was counted. Instances where mice exhibited hind paw tremor, often prior to a jump or during rearing, were also counted as paw tremor.

Sniffing: Movements of the nose and whiskers (distinct from chewing), often accompanying rearing or head-scanning behavior prior to and while walking.

**Galnon Feeding Test**

C57 Bl/6J mice were single-housed one week prior to testing. Mice had food removed 24 h before behavioral testing to increase motivation to eat. The next day, 15 min prior to the start of testing, mice were given an i.p. injection of either galnon (2 mg/kg) or vehicle. A pre-weighed pellet of standard mouse chow was introduced into the home cage at the end opposite to the mouse's location, and the latency to eat the food was timed. The test ended when the mouse bit the pellet and started consuming the food, or once 5 min elapsed, whichever occurred first. Thirty min after the start of each feeding test, the food pellet was re-weighed, and the amount consumed was calculated.
